# *C9orf72* ALS-causing mutations lead to nucleoporin Nup107 aggregation and subsequent pathological changes

**DOI:** 10.1101/2025.04.16.649118

**Authors:** Saygın Bilican, Yara Nabawi, William Hongyu Zhang, David Vilchez

## Abstract

Amyotrophic lateral sclerosis (ALS) is a fatal disorder caused by motor neuron degeneration. Hexanucleotide repeat expansions in the *C9orf72* gene, the most common genetic cause of ALS (C9-ALS), drive toxicity through different mechanisms. These pathological changes include alterations in stress granules (SGs), ribonucleoprotein complexes formed under stress conditions. Here, we show that G3BP1, a core component of SGs, aberrantly interacts with the nucleoporin Nup107 in motor neurons derived from patient iPSCs carrying *C9orf72* mutations. Moreover, Nup107 colocalizes with SGs and aggregates in C9-ALS motor neurons. Notably, knockdown of the *C. elegans* ortholog of *Nup107* alleviates ALS-associated phenotypes in worm models, including reduced lifespan and motility. Our findings provide insights into C9-ALS pathogenesis and identify Nup107 as a potential therapeutic target.

## Introduction

Amyotrophic lateral sclerosis (ALS) is the most common motor neuron disease, affecting approximately 6 in 100,000 people [1,2]. ALS leads to the degeneration of upper and lower motor neurons, resulting in muscle weakness, atrophy, and ultimately, death [3]. While the majority of ALS cases are sporadic, around 10% are familial and linked to mutations in one of over 30 different genes [3–5]. The intronic GGGGCC hexanucleotide expansion (HRE) in the first intron of *C9orf72* gene is the most common mutation causing ALS (C9-ALS), accounting for up to 40% of familial and 10% of sporadic cases [6–9].

The pathogenic mechanisms of C9-ALS are complex and non-mutually exclusive. The HRE mutation reduces *C9orf72* mRNA and protein levels, leading to a loss of function [10–12]. Additionally, GC-rich DNA and RNA repeats form G-quadruplexes that sequester nucleic acid-binding proteins, disrupting transcription and generating RNA foci [13–15]. The mutant transcript can also undergo repeat-associated non-AUG (RAN) translation, producing dipeptide repeats (DPRs) [16–19]. These DPRs exert proteotoxic effects through their aggregation and binding to low-complexity-domain containing proteins [18–28].

The toxic effects of RNA foci and DPRs in C9-ALS are closely linked to cellular stress responses, particularly stress granules (SGs) [29,30]. SGs are membrane-less organelles that form in response to different internal and external stressors but rapidly disassemble upon stress removal [31,32]. Their main components include mRNA, RNA-binding proteins (RBPs), 40S ribosomal subunits, and translation initiation factors [33–35]. Aberrant SG dynamics—such as excessive formation, aggregation, impaired disassembly, and persistence—are hallmarks of ALS [36–39]. ALS-relevant proteins such as TDP-43, FUS and DPRs, as well as HRE-RNA accumulate into SGs, establishing SGs as potential modulators of ALS pathophysiology [20,40–42]. In addition to dysregulated SG metabolism, ALS-causing *C9orf72* mutations also disrupt nucleocytoplasmic transport (NCT) pathways [43–46]. Importantly, NCT components mislocalize into SGs under stress and in ALS models [47,48]. However, the precise mechanisms linking SGs and NCT dysfunction in ALS remain unclear.

In this study, we show that the nuclear pore complex (NPC) subunit Nup107 mislocalizes, forming cytoplasmic and perinuclear foci in cells derived from C9-ALS patients. Cells carrying the *C9orf72* HRE also exhibit increased SG formation, with SGs colocalizing with Nup107 foci and displaying slower disassembly rates. Moreover, Nup107 aggregation is exacerbated under oxidative stress in C9-ALS iPSCs and their motor neuron counterparts

(iMNs). Notably, knocking down the *C. elegans* ortholog of *Nup107* in a C9-ALS model mitigates disease-associated phenotypes, including shortened lifespan, reduced motility, and DPR accumulation. Our findings establish a link between Nup107 dysregulation and altered SG dynamics, providing insights into C9-ALS pathogenesis and identifying Nup107 as a potential therapeutic target.

## Materials and Methods

### iPSC lines and culturing

The C9-1 (CS29iALS-C9n1; HRE: ∼6000-8000 bp repeat expansion) and C9-2 (CS30iALS-C9n1; HRE: ∼2400 bp repeat expansion) iPSC lines were obtained from the Cedars-Sinai RMI iPSC Core. Control and FUS^P525L^ iPSC lines were generously provided by Prof. Irene Bozzoni [49]. TDP-43^M337V^ iPSCs were obtained from RIKEN Bioresource Research Center [50]. iPSCs were maintained in mTeSR1 medium (Stem Cell Technologies,

#85850), and cultured on tissue culture-treated plates coated with Geltrex (Gibco, #A1413202) at 37°C in a humidified incubator with 5% CO₂. For passaging, cells were detached using Accutase (Life Technologies, #A1110501). Briefly, the medium was removed, and cells were incubated with Accutase for 3–5 min. Accutase was then neutralized with mTeSR1, and the cell suspension was centrifuged at 300 × *g* for 5 min. The cells were then reseeded onto Geltrex-coated plates at the desired density in mTeSR1 supplemented with 10 µM ROCK inhibitor (Abcam, # ab120129) for the first day of culture.

### Differentiation of iPSCs into motor neurons

The differentiation protocol was adapted from Hill et al., 2016 [51]. Briefly, iPSCs were cultured until they reached 80–90% confluency. The first day of media replacement was defined as day 0 (d0). From d0 to d6, mTeSR1 was replaced with Differentiation Media, cnsisting of a 1:1 mixture of DMEM F-12 (Gibco, #11320074) and Neurobasal Media (Gibco, #21103049) supplemented with 1X B27 (Gibco, #12587-010), 1X N2 (Gibco, #175020-01), 1X non-essential amino acids (Gibco, #11140050), 100 U ml^-1^ penicillin/streptomycin (Gibco, #15070063), 1X GlutaMAX (Gibco, #35050061). Additionally, the medium was supplemented with 1 μM smoothened agonist (SAG) (Sigma, #566661), 1 μM retinoic acid (RA) (Sigma, #R2625), 10 μM SB-431542 (Miltenyi Biotec, #130-106-275), and 0.1 μM LDN-193189 (Miltenyi Biotec, #130-103-925). From d7 to d13, the Differentiation Media was supplemented with 1 μM SAG, 1 μM RA, 4 μM SU-5402 (Sigma, #SML0443-5MG), and 5 μM DAPT (Sigma, #D5942) instead.

On d14, the cells were passaged using Accutase and plated onto laminin-coated tissue culture plates. Briefly, plates were first coated with 1.5 μg ml^-1^ poly-L-ornithine (L-PO) (Sigma, # P3655) in PBS and incubated overnight at 4 °C. The following day, L-PO was washed off twice with PBS and once with DMEM/F-12. Plates were then coated with 10 μg ml^-1^ mouse laminin (Gibco, #23017-015) in DMEM/F-12 and incubated overnight at 4 °C. Cells were subsequently seeded and maintained in Neuron Media (Neurobasal medium with 1X B27, 1X N2, 1X non-essential amino acids, 100 U penicillin/streptomycin, 1X GlutaMAX, 10 ng ml^-1^ BDNF [Peprotech, #450-02], and 10 ng ml^-1^ GDNF [Peprotech, #450-10]). Cells were allowed to mature for a minimum of 2 days before proceeding with subsequent experiments.

### SG induction and disassembly

SG assembly was induced by treating the cells with 500 μM sodium arsenite (Sigma, #106277) treatment. For recovery experiments, cells were washed once with DPBS (Gibco, #14200075) after the treatment and maintained in fresh culture medium. The cells were then fixed with 4% paraformaldehyde (PFA) for 15 min and stained for immunofluorescence microscopy according to the following protocol. SGs were manually counted using ImageJ [52].

### Immunofluorescence staining

Cells were plated onto glass coverslips and fixed with 4% PFA (Polysciences, #04018-1) for 15 min at room temperature, followed by two washes with 1X PBS. Cells were then permeabilized using PBS containing 0.2% (v/v) Triton X-100 and blocked with 3% bovine serum albumin (BSA) in PBS. Cells were incubated with primary antibodies diluted in 3% BSA in PBS, including mouse anti-G3BP1 (Abcam, #ab56574, 1:300), rabbit anti-NUP107 (Proteintech, #19217-1-AP, 1:50), and rabbit anti-G3BP1 (Biozol, #MBL-RN048PW, 1:500). Primary antibody incubation was carried out for 1.5 h at room temperature or overnight at 4 °C in a humidified chamber.

Following primary antibody incubation, coverslips were washed three times with 1X PBS and subsequently incubated for 45 min with secondary antibodies and 10 µg ml^-1^ Hoechst 33342 (Invitrogen, #H3570), all diluted in 3% BSA in PBS. The secondary antibodies used were Goat anti-Mouse IgG (H+L) Highly Cross-Adsorbed Secondary Antibody, Alexa Fluor 488 (Thermo Fisher, #A-11029, 1:300) and Goat anti-Rabbit IgG (H+L) Cross-Adsorbed Secondary Antibody, Alexa Fluor 594 (Thermo Fisher, #A-11012, 1:300). Unbound secondary antibodies were removed by three washes with PBS, followed by a final rinse with ddH₂O. Coverslips were dehydrated using 100% ethanol and left to dry in the dark. Once dried, coverslips were mounted onto microscope slides using ProLong Diamond Antifade Mountant (Invitrogen, #P36961) or FluorSave Reagent (Merck, #345789). Imaging was performed using a Zeiss Axio Imager Z.1 microscope.

### Western blotting

Protein extraction was performed using either RIPA buffer (50 mM Tris-Cl pH 7.5, 150 mM NaCl, 1% Triton X-100, 1% sodium deoxycholate, 0.1% SDS, 1 mM EDTA) or a non-denaturing native lysis buffer (150 mM NaCl, 50 mM HEPES pH 7.4, 1 mM EDTA and 1% Triton X-100). Both buffers were supplemented with cOmplete Mini Protease Inhibitor Cocktail (Roche, #11836153001) and 1 mM PMSF. Equal amounts of extracted protein were separated via SDS-PAGE and transferred onto PVDF membranes. Membranes were blocked with 3% BSA in TBS-T for 1 h at room temperature, followed by overnight incubation at 4 °C with primary antibodies diluted in blocking solution (mouse anti-poly-GA [Merck, #MABN889, 1:1,000], mouse anti-α-tubulin [Sigma, #T6199, 1:5,000], rabbit anti-NUP107 [Proteintech, #19217-1-AP, 1:5,000]). The next day, membranes were washed three times with TBS-T (5 minutes per wash) and incubated for 1 h at room temperature with secondary antibodies diluted in blocking solution (Donkey HRP AP anti-Mouse IgG (H+L) [Jackson Immuno Research, #715-035-1500, 1:10,000], Donkey HRP AP anti-Rabbit IgG (H+L) [Jackson Immuno Research, #715-035-1520, 1:10,000]). Following secondary antibody incubation, membranes were washed three times with TBS-T (5 min per wash) and developed using Immobilon Western Chemiluminescent HRP Substrate (Merck, #WBKLS0500). The signal was detected using the FUSION SOLO S imaging system (Vilber).

### Filter trap assays

Proteins were extracted using native lysis buffer, and lysates were sonicated for 30 s at 40% amplitude using the Bandelin Electronic Sonopuls Ultrasonic Homogenizer Mini20. Cell debris was removed by centrifugation at 3,000 *g* for 4 min at 4 °C, and the supernatant was transferred to a new tube. Equal protein amounts from each sample were adjusted to 100 µL and supplemented with a final concentration of 0.5% SDS. The slot blot apparatus (BIO-RAD, #1706542) was assembled with a cellulose acetate membrane (VWR, #516-5020) and equilibrated with native buffer containing 0.5% SDS. Samples were loaded into the slots and allowed to pass through the membrane completely. The membrane was then washed with 0.2% SDS in ddH₂O.

The membrane was then blocked with 3% BSA in TBS-T for 1 h and incubated overnight at 4 °C with the primary Nup107 antibody (rabbit anti-NUP107 [Proteintech, #19217-1-AP, 1:5,000]). The next day, the membrane was washed three times with TBS-T and incubated with the secondary antibody IRDye 800CW Donkey anti-Rabbit IgG (H + L) (LI-COR, #926-32213, 1:10,000) for 1 h. After incubation, the membrane was washed again with TBS-T, and imaging was performed using the Odyssey M Imaging System.

### Co-immunoprecipitation and proteomics sample preparation

Proteins were extracted from neuronal cultures using SDS-free RIPA buffer and homogenized with a 27-gauge needle. After homogenization, the samples were centrifuged at maximum speed for 10 min at 4 °C, and the resulting supernatants were collected. For immunoprecipitation, 300 µg of protein lysates were incubated with 2 µg of monoclonal mouse anti-G3BP1 antibodies (Abcam, #ab56574) on ice for 1 h. Next, 50 µL of Protein A microbeads (Miltenyi Biotec, #130-071-001) were added, and the mixture was incubated at 4°C on a rotator for 2 h. All subsequent steps were performed in a cold room until the digestion step.

µColumns (130-042-701, Miltenyi Biotec) placed on magnetic holders were pretreated with 200 µL of SDS-free RIPA buffer. The protein-antibody-bead complex was loaded onto the columns and allowed to pass through by gravity flow. The columns were washed three times with 200 µL of Wash Buffer I (50 mM Tris-Cl pH 7.5, 150 mM NaCl, 0.05% Triton X-100, 5% glycerol), followed by five washes with Wash Buffer II (50 mM Tris-Cl pH 7.5, 150 mM NaCl).

Bound proteins were digested directly on the columns with 25 µL digestion buffer (2 M Urea, 7.5 mM ammonium bicarbonate, 1 mM DTT, 5 ng ml^-1^ trypsin) at room temperature for 30 min, then eluted using 50 µL of elution buffer (2 M Urea, 7.5 mM ammonium bicarbonate, 5 mM CAA). Digestion was completed overnight in the dark. The next day, formic acid was added to a final concentration of 4%, and samples were centrifuged at maximum speed for 5 min. The supernatants were loaded onto SDP-RP StageTips and sent for label-free quantitative proteomics at the CECAD Proteomics Core Facility. Raw proteomics data were analyzed using Perseus v1.6.13.0 [53].

### *C. elegans* strains and maintenance

*C. elegans* were maintained under standard conditions at 20°C on nematode growth medium (NGM) plates seeded with the OP50 *E. coli*. The KRA315 (*snb-1p*::*C9^ubi^* + *myo-2p*::GFP) and KRA317 (*snb-1p*::*ΔC9^ubi^* + *myo-2p*::GFP) strains were kindly provided by Paschalis Kratsios [54]. For RNAi experiments, late-L4 stage *C. elegans* were fed with the *E. coli* HTT115 strain carrying either the L4440 empty vector control or L4440 expressing double-stranded RNA targeting *npp-5*. The *npp-5* RNAi construct was obtained from the Ahringer library and verified by sequencing using the forward primer 5’– GTAAAACGACGGCCAGTGAG–3’.

### *C. elegans* lifespan assays

Synchronization of *C. elegans* was carried out using a bleaching protocol. Adult hermaphrodites containing eggs were treated with a bleaching solution (1.5% v/v NaClO, 0.75 M KOH in ddH_2_O) until the eggs were released and the adults were dissolved. The eggs were then incubated overnight in M9 buffer to allow hatching and synchronization at the L1 larval stage. L1 larvae were transferred to OP50-seeded plates and grown at 20°C until they reached the first day of adulthood.

For lifespan assays, 96 worms per condition were transferred onto RNAi plates and monitored either daily or every other day [55]. Worms exhibiting a protruding vulva, bagging phenotype, or that were lost during the experiment, were censored. Lifespan data were analyzed using GraphPad Prism version 6.0 and statistical significance was calculated by the log-rank (Mantel-Cox) test. OASIS software was used to determine mean lifespan. P-values were calculated for comparisons between two groups within a single experiment.

### *C. elegans* motility assays

Day-5 adult *C. elegans* were placed in 20 µL of M9 buffer and allowed to acclimate for 30 s. Body bends were defined as changes in body direction. The number of body bends was counted over the next 30 s [56].

### Bacterial expression and purification of G3BP1 and Nup107

The protocol for protein expression and purification was adapted from Llamas et. al 2023 [57]. Briefly, human G3BP1 and NUP107 cDNA were cloned into pGEX-6P-1 vector, which contains a 6x His-Tag, using GeneArt Gibson Assembly HiFi Master Mix (ThermoFischer, #A46627) and the primers listed in **Supplementary Data 1**. After confirming the constructs by sequencing, *E. coli* BL21(DE3) were transformed with the vector containing the respective cDNA. The bacteria were cultured at 37 °C for initial growth, and the culturewas further incubated at 18 °C overnight after the addition of 0.25 mM isopropyl 1-thio-β-d-galactopyranoside to induce protein expression.

The bacterial culture was centrifuged at 25,000 *g* at 4 °C for 1 h and then sonicated. The lysates were clarified by centrifugation at 15,000 g in 4 °C for 1 h. His-Tag containing proteins were purified using HisPurCobalt Resin (Thermo Scientific, #89964) via affinity chromatography. The His-tag was removed by treatment with TEV protease during overnight dialysis. The dialyzed proteins were subjected to a second affinity chromatography step to remove contaminants, and enriched fractions were concentrated using Amicon Ultra-15 filters (Merck, #10403892). Protein concentration was determined with NanoDrop 8000. Single-use aliquots of the purified proteins were snap-frozen in liquid nitrogen and stored at −80 °C. Each protein fraction obtained during the purification process was analyzed by SDS-PAGE.

### Electrophoretic Mobility Shift Assay (EMSA)

The EMSA protocol was adapted from Hsieh et. al 2016 and Celona et. al 2017 [58,59]. The (GGGGCC)_6.5_ probe, tagged with TYE 563 tag at its 5’end, was obtained from IDT. Reaction mixtures containing 0.5 µM of the probe and varying concentrations of the protein of interest were prepared in protein-RNA binding buffer (40 mM Tris-Cl pH 8, 30 mM KCl, 1 mM MgCl_2_, 0.01% (v/v) Nonidet P40 Substitute, 1 mM DTT, 5% (v/v) glycerol, 10 µg ml^-1^ BSA), adjusted to a final volume of 20 µL. The mixtures were incubated at room temperature for 30 min in the dark.

The native gels for EMSA assays (5% polyacrylamide, 0.5x TBE and 2.5% glycerol) were pre-run in 0.5x TBE (45 mM Tris, 45 mM boric acid, 1 mM EDTA pH 8) at 100V for at least 30 min before loading the samples. After the pre-run, samples were mixed with Orange G loading dye and run at 100V until the loading dye reached the bottom of the gel. Imaging was performed using the Odyssey M Imaging System.

### Statistical Analysis

Statistical analyses were performed using GraphPad Prism (versions 9 and 10). The specific statistical tests used and their corresponding significance levels are described in the figure legends.

## Results

Cumulative evidence suggests that SGs act as a nidus for pathological protein aggregation, particularly when they lose their dynamic properties and transition into persistent granules [38,40,60,61]. In line with this, ALS-causing mutations, including abnormal hexanucleotide expansions in *C9orf72*, induce SG alterations [21,22,28,29,40,49,62–66]. To gain insights into the pathogenic interplay between SGs and ALS, we assessed SG dynamics in two patient-derived iPSC lines carrying distinct *C9orf72*-HRE mutations (C9-1: ∼6000-8000 bp repeat expansion; C9-2: ∼2400 bp repeat expansion). To this end, we monitored SG assembly and disassembly at various time points during arsenite treatment (oxidative stress) and after stress removal (**Fig. 1A-B**).

**Figure 1.**
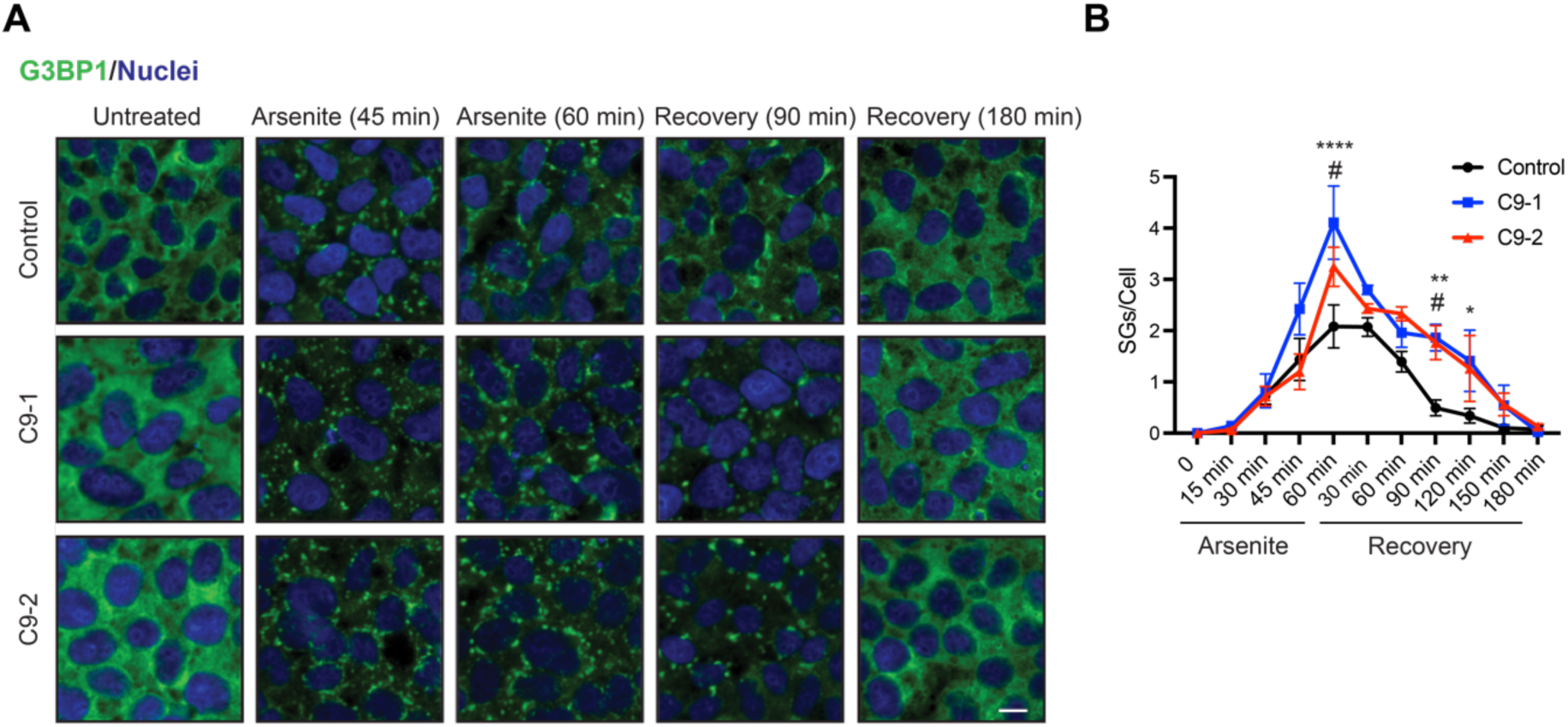
ALS-causing *C9orf72* mutations induce alterations in SG dynamics. (**A**) Immunocytochemistry in control and C9-ALS iPSCs during treatment with 500 µM arsenite and at the indicated time points following arsenite removal (recovery). G3BP1 and Hoechst 33342 staining were used as markers of SGs and nuclei, respectively. Images are representative of three independent experiments. Scale bar: 10 μm. (**B**) Quantification of total G3BP1-positive SGs per cell in iPSC cultures (mean ± s.e.m.). Data were obtained from three independent experiments, with at least 200 cells counted per condition. Statistical comparisons were made by two-way ANOVA with Šidák multiple-comparison test. *P* values: * C9-1 vs control, *P* < 0.05; ** C9-1 vs control *P* < 0.01; **** C9-1 vs control, *P* < 0.0001; ^#^ C9-2 vs control, *P* < 0.05.

Both C9-ALS mutant lines (C9-1 and C9-2) exhibited an increased number of SGs after 1 hour of arsenite treatment compared to controls. Notably, C9-1 cells, which harbor a longer HRE, displayed elevated SG numbers upon arsenite treatment, compared to C9-2 (**Fig. 1A-B**). During recovery, both mutant cell lines exhibited slower SG disassembly than controls (**Fig. 1A-B**). This trend persisted until 3 hours post-recovery, at which point SG disassembly was complete in both control and mutant *C9orf72*-expressing lines (**Fig. 1A-B**).

Given that SG composition influences their dynamics, we analyzed changes in the interactome of the SG core protein G3BP1 in iPSC-derived motor neurons (iMNs). To this end, we performed co-immunoprecipitation of G3BP1 followed by label-free proteomics under both basal and arsenite-induced SG assembly. Upon SG assembly, we identified 40 proteins with increased interaction with G3BP1 in both C9-1 and C9-2 mutant lines compared to control motor neurons (**Fig. 2A and Supplementary Data 2**). Among them, we detected several neurodegeneration-relevant proteins, such as HSPH1 [67], UBA1 [68], DDX6 [69], SEPT9 [70], and Nup107 [45] (**Fig. 2A and Supplementary Data 2**). Importantly, the interaction between G3BP1 and Nup107 was significantly upregulated in mutant *C9orf72* motor neurons upon SG formation following arsenite treatment (**Fig. 2B and Supplementary Data 2**). We focused on Nup107 due to its critical role in nucleocytoplasmic transport, a pathway widely implicated in C9-ALS, although the specific contributions of individual components remain unclear [43–45,71–73]. Furthermore, Nup107 was the only subunit of the nuclear pore complex (NPC) found to be enriched in the SGs of mutant *C9orf72* motor neurons (**Supplementary Data 2**).

**Figure 2.**
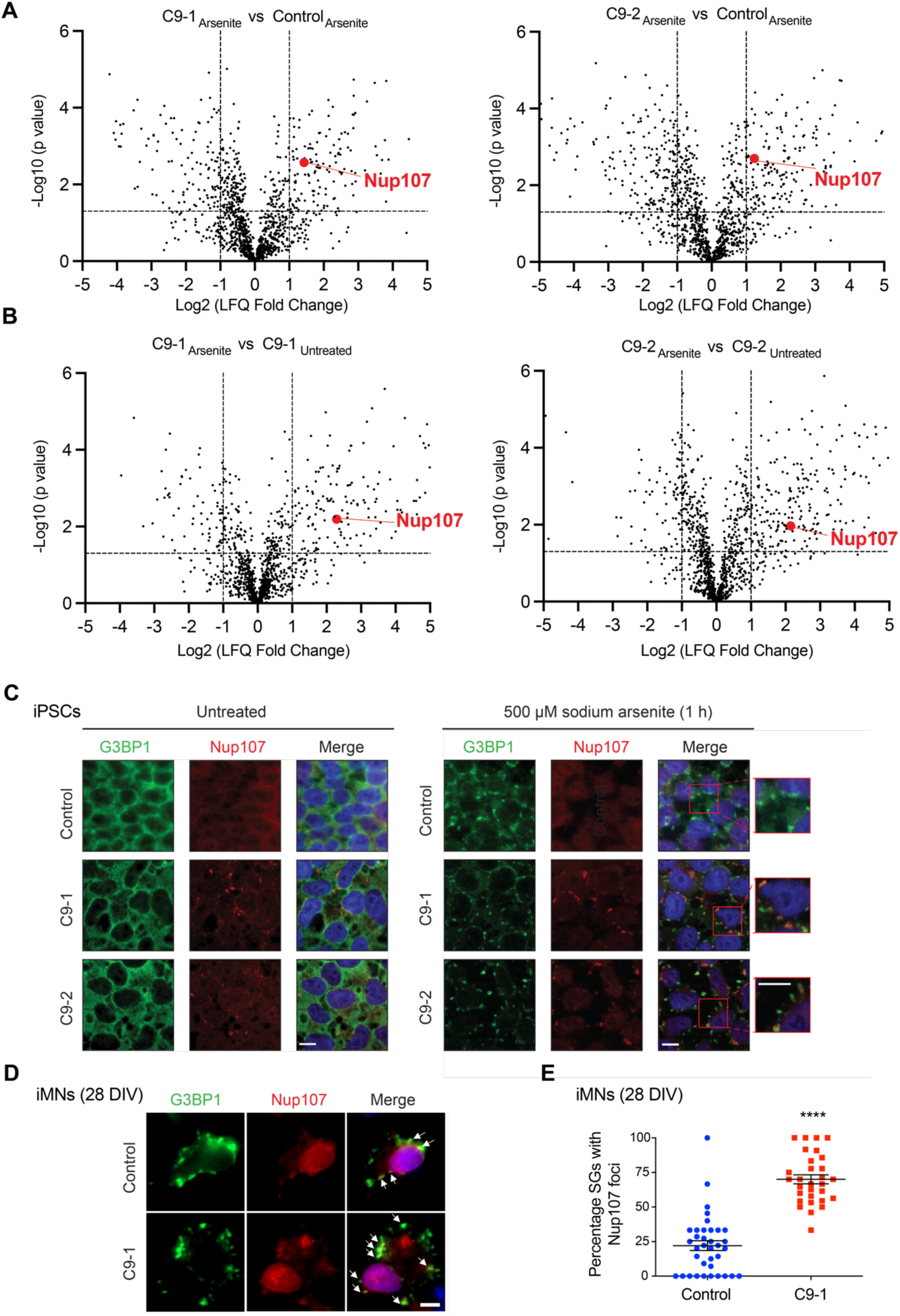
G3BP1 gains enhanced interaction with Nup107 in mutant *C9orf72* motor neurons upon SG assembly. (**A**) Volcano plots of the G3BP1 interactome in *C9orf72* motor neurons (C9-1: ∼6000-8000 bp repeat expansion; C9-2: ∼2400 bp repeat expansion) compared to control motor neurons. iPSC-derived motor neurons (iMNs) from C9-ALS patients and controls were treated with 500 μM sodium arsenite for 1 h. The −log_10_(*P* value) of a two-sided *t*-test is plotted against the log_2_-transformed fold change of protein label-free quantification (LFQ) values from immunoprecipitation with anti-G3BP1 antibody (*n* = 3 biological replicates). (**B**) Volcano plots comparing the G3BP1 interactome in *C9orf72* motor neurons with and without 500 μM sodium arsenite treatment (1 h) (*n* = 3 biological replicates). (**C**) Immunocytochemistry using G3BP1 and Nup107 antibodies in control and C9-ALS iPSCs under basal conditions (untreated) or after treatment with 500 µM sodium arsenite for 1 hour. Hoechst 33342 staining (blue) was used to label nuclei. Images are representative of three independent experiments. Scale bar: 10 μm. (**D**) Immunocytochemistry with G3BP1 and Nup107 antibodies in iMNs treated with 500 µM sodium arsenite (1 h) after 28 days in vitro (DIV). Hoechst 33342 staining (blue) was used as a marker of nuclei. Images are representative of two independent experiments. Arrow indicates colocalization of SGs with Nup107 foci. Scale bar: 5 μm. (**E**) Percentage of SGs containing Nup107 foci in iMNs (28 DIV) treated with 500 µM sodium arsenite for 1 hour (mean ± s.e.m.; Control *n* = 36 neurons; C9-1 *n* = 30 neurons from 2 independent experiments). Statistical comparison was made by two-tailed Student’s *t*-test for unpaired samples (**** *P* < 0.0001).

To validate our proteomics findings, we performed immunostaining experiments using antibodies against G3BP1 and Nup107. In undifferentiated control iPSCs, Nup107 exhibited a cytoplasmic and perinuclear distribution under both basal conditions and arsenite treatment (**Fig. 2C**). In contrast, C9-ALS iPSCs displayed cytoplasmic Nup107 foci even in the absence of stress (**Fig. 2C**). Upon arsenite-induced stress, these Nup107 foci co-localized with SGs (**Fig. 2C**). Following differentiation into motor neurons, iMNs treated with arsenite after two days in vitro did not show co-localization of Nup107 with SGs (**Supplementary Fig. 1**). However, when motor neurons were treated with arsenite after 28 days in vitro, we observed a significant increase in the percentage of SGs containing Nup107 foci in C9-ALS iMNs compared to controls (**Fig. 2D–E**). These results suggest that mutant *C9orf72*–induced mislocalization of Nup107 into SGs may worsen over time in motor neurons.

Nup107 is a component of the Y-complex, a critical subcomplex required for NPC assembly [74]. To determine whether other structurally essential NPC subunits exhibit altered intracellular distribution in C9-ALS cells, we screened additional components of the Y-complex. Besides Nup107, none of the other five tested Y-complex proteins showed SG colocalization or changes in intracellular distribution in C9-ALS cells compared to controls (**Supplementary Fig. 2A-E**). Additionally, to assess whether Nup107 mislocalization is specific to C9-ALS, we examined cells expressing ALS-related FUS^P525L^ and TDP-43^M337V^ mutant variants. However, mutant FUS and TDP-43 cells did not exhibit altered Nup107 distribution or colocalization with SGs (**Supplementary Fig. 3**).

Protein aggregation is a hallmark of ALS pathology, and certain nucleoporins can aggregate with molecular crowders or during aging, contributing to proteotoxic stress [75–77]. Although Nup107 lacks the phenylalanine-glycine repeats typically associated with nucleoporin aggregation, its intrinsically disordered region may promote aggregation (**Supplementary Fig. 4**). To determine whether Nup107 aggregates in C9-ALS cells, we performed filter trap experiments to assess SDS-insoluble protein aggregates. Under normal conditions, we did not detect insoluble Nup107 aggregates in undifferentiated C9-ALS iPSCs (**Fig. 3A**). However, arsenite treatment led to the accumulation of Nup107 aggregates in these cells (**Fig. 3A**). In contrast, control iPSCs did not exhibit Nup107 aggregation upon SG assembly following arsenite treatment, despite similar total Nup107 levels between ALS and control lines (**Fig. 3A**). Given that motor neurons are the primary cell type affected in ALS, we next examined iMNs. Notably, C9-ALS iMNs displayed elevated Nup107 aggregation even in the absence of arsenite treatment (**Fig. 3B**).

**Figure 3.**
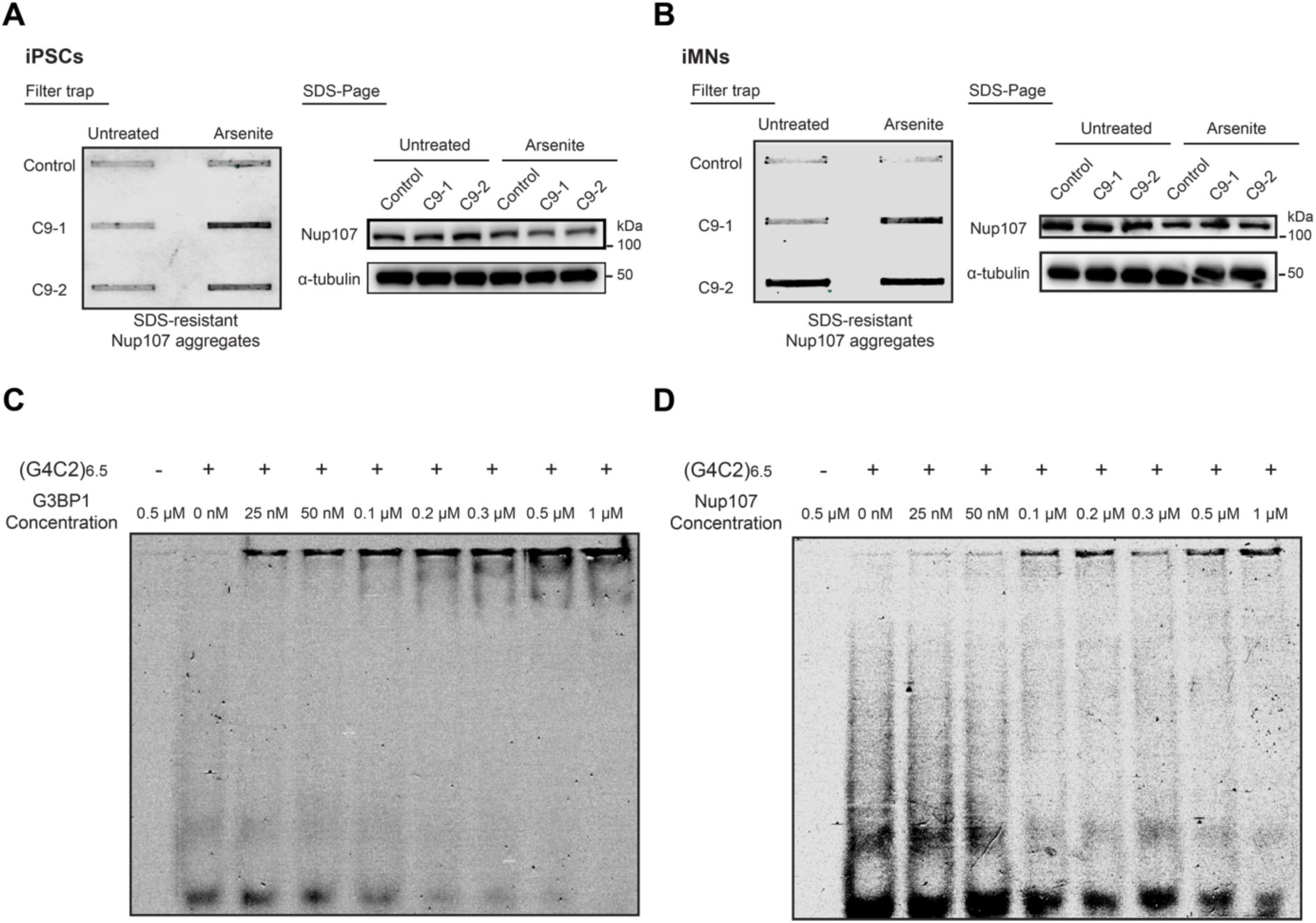
Nup107 aggregates in C9-ALS cells and binds to pathological *G_4_C_2_* RNA repeats. (**A**) Filter trap assay detecting Nup107 aggregation (anti-Nup107 antibody) in control and C9-ALS iPSCs under basal conditions (untreated) and following treatment with 500 µM arsenite for 1 hour. Right: SDS–polyacrylamide gel electrophoresis (SDS-PAGE) with antibodies to Nup107 and α-tubulin loading control. Images are representative of three independent experiments. (**B**) Filter trap of Nup107 aggregates in control and C9-ALS iMNs under basal conditions and after 1 hour of 500 µM sodium arsenite treatment. Right: SDS-PAGE with antibodies to Nup107 and α-tubulin. Images are representative of three independent experiments. (**C-D**) Electrophoretic mobility shift assays (EMSA) using purified recombinant G3BP1 (**C**) or Nup107 (**D**) titrated with fluorescently labeled (*G4C2*)₆.₅ RNA. The presence of the probe and protein concentrations are indicated at the top. Images are representative of two independent experiments.

Prompted by these findings, we investigated whether HRE-derived *G_4_C_2_* RNA repeats directly interact with G3BP1 and Nup107 in C9-ALS cells, as RNA-protein interactions can change protein solubility [78–81]. To assess binding affinity, we performed electrophoretic mobility shift assays (EMSA) by titrating purified recombinant G3BP1 and Nup107 with fluorescently labeled (*G4C2*)_6.5_ RNA (**Fig. 3C-D**). Even at low concentrations (25 nM), G3BP1 bound repeat RNA, as indicated by increased retention of the RNA-protein complex in the gel well (**Fig. 3C**). This binding was dose-dependent, with increasing G3BP1 levels leading to a reduction in free RNA at the bottom of the gel (**Fig. 3C**). On the other hand, Nup107 exhibited weaker binding affinity, with detectable binding only at 100 nM and no further increase at higher concentrations (**Fig. 3D**). These findings suggest that pathological RNA repeats directly interact with Nup107 and G3BP1, potentially contributing to their aggregation.

Given the links between Nup107 and C9-ALS, we asked whether Nup107 influences pathological phenotypes *in vivo*. To address this, we used a previously established *C. elegans* model of C9-ALS that expresses 75 repeats of the *G_4_C_2_* HRE, flanked by *C9orf72* intronic sequences and driven by the ubiquitous *snb-1* promoter (*C9^ubi^*) [54] (**Fig. 4A**). As a control, we used worms expressing the same construct without HRE (*ΔC9^ubi^*) (**Fig. 4A**). *C9^ubi^* worms exhibit key C9-ALS pathophysiological features, including DPR production via RAN translation, progressive motility decline, and reduced lifespan compared to *ΔC9^ubi^* worms [54]. To investigate the role of Nup107 in ALS pathology, we knocked down its *C. elegans* ortholog, *npp-5*, which shares 43% coverage and 41% sequence similarity with human Nup107, as determined by BLAST analysis [82]. To achieve this, worms were fed RNAi bacteria expressing either an empty vector control or *npp-5*-targeting RNAi after development.

**Figure 4.**
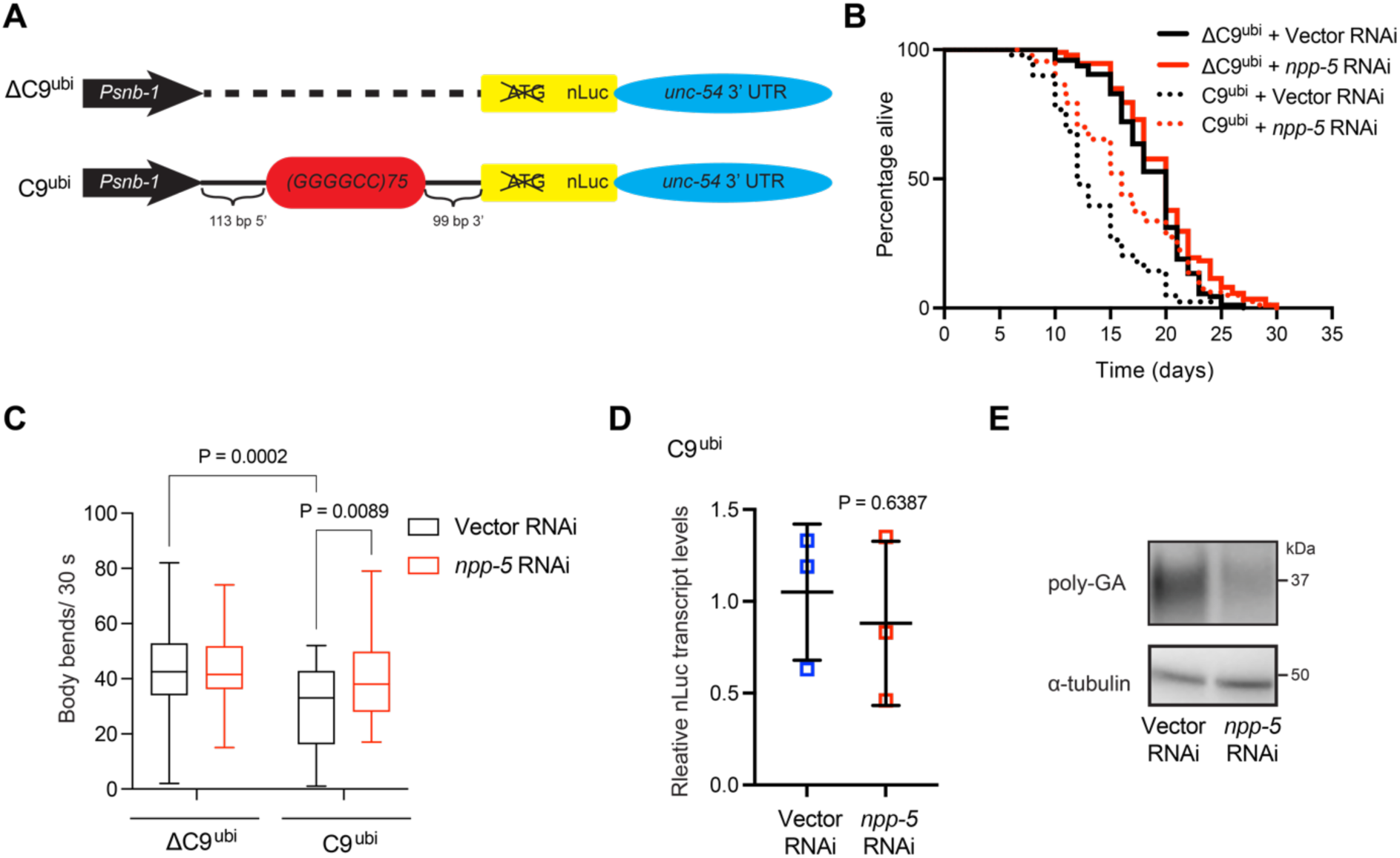
Knockdown of *npp-5*, the *C. elegans Nup107* ortholog, ameliorates disease-related phenotypes in C9-ALS worm models. (**A**) Schematic representation of transgenic constructs ΔC9^ubi^ and C9^ubi^ for C9-ALS modeling in *C. elegans*. The schematic is adapted from the original publication describing the generation of these strains (Sonobe et. al 2021 [54]). (**B**) ALS *C9^ubi^* worms have a shorter lifespan compared to control *ΔC9^ubi^* worms (*P* <0.0001). Knockdown of *npp-5* after development alleviates the short lifespan phenotype of ALS *C9^ubi^* worms (*P* <0.0001). *ΔC9^ubi^* + vector RNAi mean ± s.e.m.: 18.66 days ± 0.37; *ΔC9^ubi^* + *npp-5* RNAi: 19.72 ± 0.42; *C9^ubi^* + vector RNAi: 13.55 ± 0.44; *C9^ubi^* + *npp-5* RNAi: 16.48 ± 0.57. *P*-values: two-sided log-rank test, n= 96 worms/condition. **Supplementary Data 3** contains statistical analysis and replicate data from independent lifespan experiments. (**C**) Body bends over a 30-s period in control and C9-ALS worms at day 5 of adulthood (*n* = 40 worms per condition). Box plots represent the 25^th^–75^th^ percentiles, the lines depict the median and the whiskers show the minimum–maximum values. Statistical comparisons were made by two-way ANOVA with Fisher’s LSD test. (**D**) qPCR analysis of *nLuc* transcript levels in ALS *C9^ubi^* worms at day 5 of adulthood (relative expression to Vector RNAi, mean ± SD, *n* = 3 biological replicates). Statistical comparisons were made by two-tailed unpaired Student’s t-test. (**E**) Western blot analysis of poly-GA levels in *C9^ubi^* worms at day 5 of adulthood. α-tubulin is the loading control. Images are representative of 4 independent experiments. In all experiments, RNAi treatment was initiated after development.

Notably, knockdown of *npp-5* ameliorated the short lifespan phenotype of *C9^ubi^* worms (**Fig. 4B and Supplementary Data 3**), suggesting an improvement in disease pathology. Given that motor dysfunction is a hallmark of ALS, we assessed motility in these worms. ALS *C9^ubi^* worms exhibited a significant decline in motility compared with *ΔC9^ubi^* controls at day 5 of adulthood, a phenotype rescued by *npp-5* knockdown (**Fig. 4C**). To determine whether this phenotypic improvement was associated with changes in the transcription of the *G_4_C_2_* HRE construct or its translation into DPRs, we examined the expression of poly-GA, one of the most pathological HRE-derived DPRs. The *C9^ubi^* construct includes a nanoluciferase (*nLuc*) tag placed in the poly-GA reading frame (**Fig. 4A**). While the transcript levels of *nLuc* remained unchanged upon *npp-5* knockdown (**Fig. 4D**), western blot analysis revealed a marked reduction in poly-GA peptide levels (**Fig. 4E**). These findings suggest that lowering NPP-5/Nup107 levels does not affect HRE transcription but instead decreases DPR accumulation from HRE-derived transcripts.

## Discussion

Dysregulation of SG dynamics and nucleocytoplasmic transport (NCT) are emerging hallmarks of C9-ALS; however, the precise link between these processes remains unclear [29,45,47,62,73]. Our study identifies the nucleoporin Nup107 as a potential factor bridging these two disease-related changes. We demonstrate that Nup107 is widely mislocalized, associates with SGs, and aggregates in C9-ALS cells. Notably, our experiments reveal that Nup107 aggregation occurs independently of its total protein levels, suggesting that its accumulation into insoluble aggregates results from misfolding or sequestration rather than protein overexpression. Nup62, another nucleoporin with intrinsically disordered regions, has been shown to colocalize with G3BP1 and TDP-43 in the cytoplasm, contributing to their insolubility [48].

The impact of HRE-derived RNA on nucleoporin dysfunction has been largely unexplored. Our data reveal that Nup107 directly binds *G_4_C_2_* repeat RNA, albeit with lower affinity than G3BP1. This suggests that HRE RNA may contribute to Nup107 misfolding or altered function, potentially linking RNA toxicity to SG alterations. Along these lines, previous studies have shown that G3BP1 colocalizes with HRE sense- and antisense probes [42,83]. Furthermore, Zfp106, another HRE-interacting protein, has been reported to bind Nup107 [59], reinforcing the notion that HRE RNA contributes to nucleoporin dysfunction in C9-ALS.

To explore the role of Nup107 in ALS pathology, we examined whether reducing its levels mitigates C9-ALS phenotypes using *C. elegans* models. Knockdown of *npp-5*, the *C. elegans* ortholog of *Nup107*, significantly extended lifespan and rescued motor deficits in C9-ALS worms. These beneficial effects occurred without changes in mutant transcript levels. Instead, we observed a marked reduction in poly-GA peptide levels, suggesting that Nup107 may influence RAN-mediated translation into DPRs. Alternatively, Nup107 could influence DPR clearance, reducing its accumulation.

In summary, our findings highlight Nup107 as a potential therapeutic target in C9-ALS, linking RNA toxicity, nucleoporin dysfunction, SG alterations and protein aggregation. Indeed, our *in vivo* results suggest that reducing NPP-5/Nup107 levels mitigates ALS-related changes in worm models. These findings support the role of nucleoporins as contributors to ALS pathology and open new avenues for therapeutic intervention.

## Acknowledgements

This work was supported by the Deutsche Forschungsgemeinschaft (CRC1678 and Germany’s Excellence Strategy-CECAD, EXC 2030-390661388) and the Else Kröner-Fresenius-Stiftung (2021-EKSE.95). We thank the CECAD Proteomics Core Facility for contribution and advice on proteomics experiments.

## Abbreviations

ALS: amyotrophic lateral sclerosis
C9-ALS: *C9orf72*-amyotrophic lateral sclerosis
DPR: dipeptide repeats
EMSA: electrophoretic mobility shift assay
HRE: hexanucleotide repeat expansion
iMN: iPSC-derived motor neuron
iPSC: induced pluripotent stem cell
NCT: nucleocytoplasmic transport
NPC: nuclear pore complex
RAN: repeat-associated non-AUG translation
RBP: RNA-binding proteins
SG: stress granule

