## Supplementary Figures 1-4 for "*C9orf72* ALS-causing mutations lead to nucleoporin Nup107 aggregation and subsequent pathological changes"

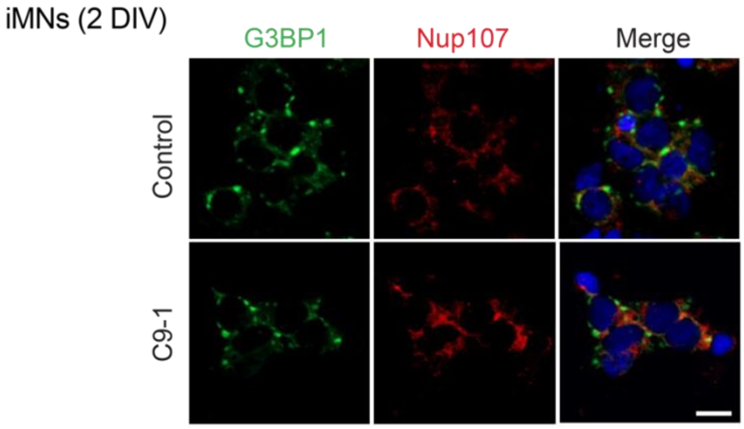


**Supplementary Figure 1. Co-localization of Nup107 foci with SGs is not observed in early iMNs.** Immunocytochemistry with G3BP1 and Nup107 antibodies in iMNs treated with 500 µM sodium arsenite (1 h) after 2 days in vitro (DIV). Hoechst 33342 staining (blue) was used as a marker of nuclei. Images are representative of two independent experiments. Scale bar: 10 µm.

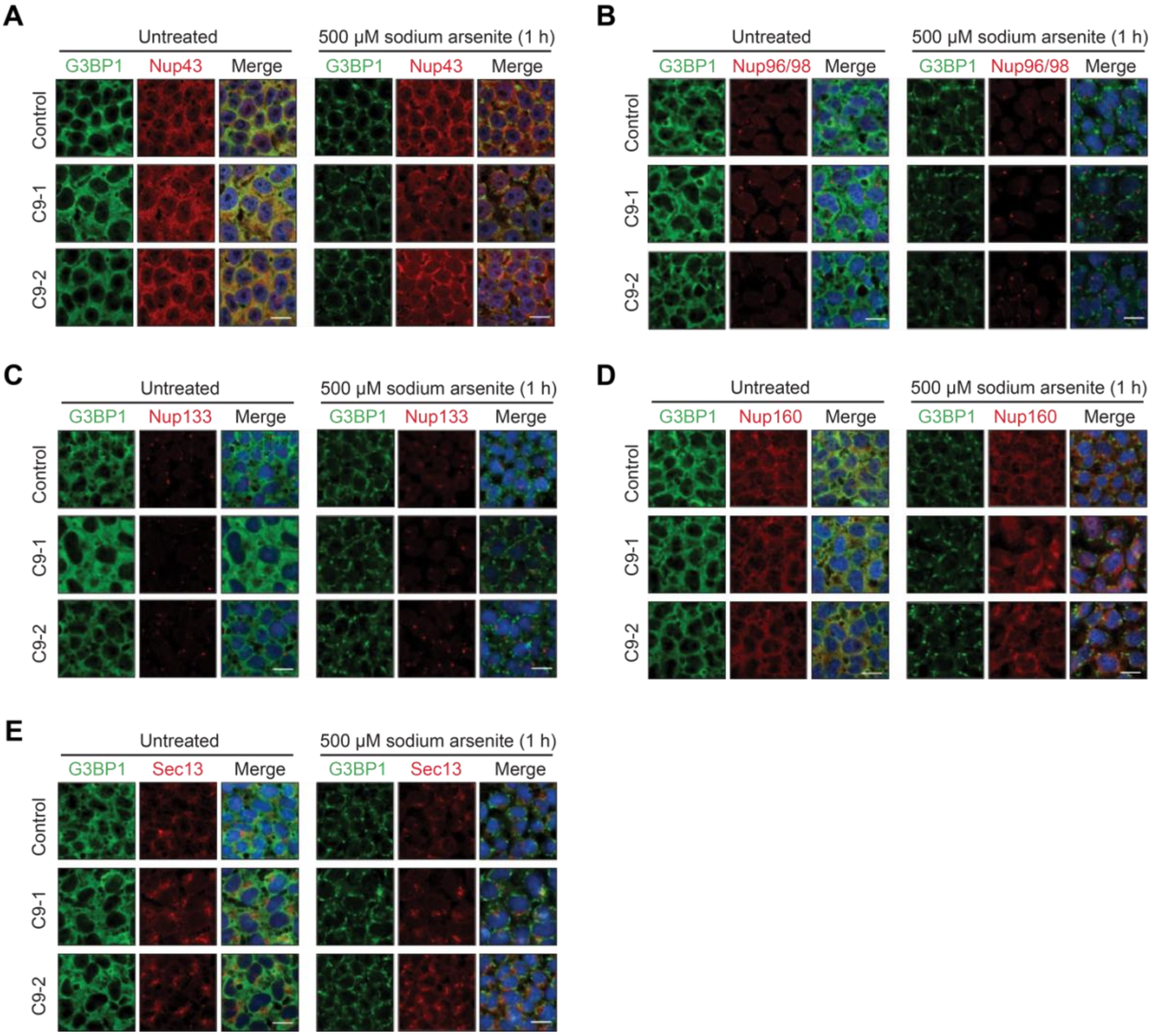


**Supplementary Figure 2. ALS-causing *C9orf72* mutations do not induce co-localization of other outer ring nucleoporins with SGs**. Immunostaining of G3BP1 and distinct Y-complex nucleoporins (Nup43 (**A**), Nup96/98 (**B**), Nup133 (**C**), Nup160 (**D**) and Sec13 (**E**)) in control and C9-ALS iPSCs under basal conditions (untreated) or after treatment with 500 µM sodium arsenite for 1 hour. Hoechst 33342 staining was used to label nuclei. Images are representative of three independent experiments. Scale bars: 20 µm.

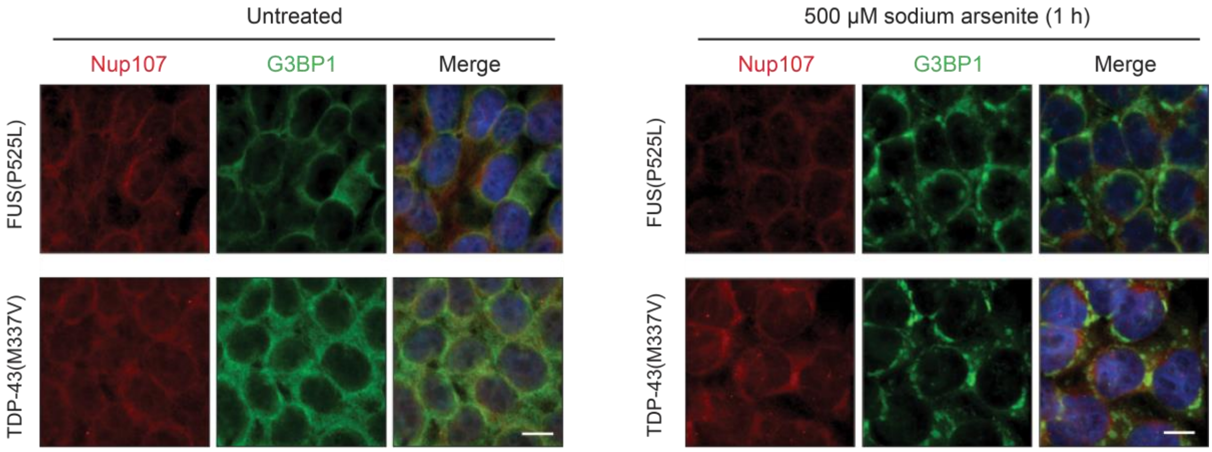


**Supplementary Figure 3. ALS-causing FUS and TDP-43 mutations do not induce NUP107 localization within SGs.** Immunostaining of iPSCs expressing ALS-related mutant FUS and TDP-43 variants after treatment with 500 µM sodium arsenite for 1 hour. Hoechst 33342 staining was used to label nuclei. Images are representative of three independent experiments. Scale bars: 10 µm.

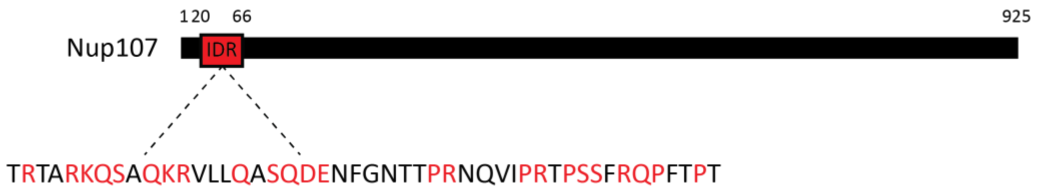


**Supplementary Figure 4. Schematic representation of the intrinsically disordered region of the Nup107 protein.**
